# The time course of co-speech gesture production: An MEG study

**DOI:** 10.64898/2026.05.04.722691

**Authors:** Kazuki Sekine, Reiji Ohkuma, Hiroshi Ban

## Abstract

People frequently gesture while speaking, even when listeners cannot see them—for instance, during phone calls or behind barriers. Congenitally blind individuals also gesture, indicating that gestures serve functions beyond visual communication. Previous models of gesture production (e.g., Kita & Özyürek, 2003; Rauscher et al., 1996) suggest that gestures facilitate speech, but they rely heavily on behavioural data and provide limited insight into temporal dynamics. This study used magnetoencephalography (MEG), a neuroimaging technique with high temporal resolution, to investigate when gestures influence speech. Twenty-three native Japanese speakers took part in a storytelling task under two conditions: Gesture-Required (gesture use instructed) and Gesture-Prohibited (hands kept still). Participants described cartoon clips across multiple sessions (30 trials × 3 sessions per condition). Using speech onset as the reference point, we compared root mean square (RMS) values within a –0.25 to 0 second window. RMS values were higher in the Gesture-Prohibited condition, with increased activity in the bilateral anterior temporal lobes (Left ATL: *p* = .049; Right ATL: *p* = .027), but not in motor regions (*p* = .29). These findings suggest that gestures reduce neural load in language-related regions before articulation. Co-speech gestures may support speech planning by facilitating lexical retrieval or semantic structuring. The lack of motor region effects indicates that this influence is linguistic rather than motoric. This study provides direct direct neurophysiological evidence of the timing of gesture–speech interaction, supporting models that view gestures as an integral part of speech production.

## 1. Background

When we speak, we often use hand gestures. These gestures, known as co-speech gestures, are spontaneously produced alongside speech (McNeill, 1992). Co-speech gestures are characterised by an idiosyncratic relationship between their form and meaning, which varies across individuals and contexts, and by their close temporal synchrony with the speech segments that convey the same or related referents (McNeill, 2005). As a result, co-speech gestures are distinct from emblems, which are gestures with form–meaning relationships determined by social conventions and which can be understood independently of speech.

They also differ from sign languages, which are fully developed linguistic systems. In what follows, we refer to co-speech gestures simply as gestures. It has been proposed that gestures not only convey communicative content but also facilitate the process of speech production (Kita, 2000). In the present study, we investigated when and how gestures influence speech during production by measuring brain activity while participants were speaking and gesturing.

Since McNeill (1992, 2005) argued that gesture and speech form a tightly integrated system in both comprehension and production, research has increasingly shifted away from the traditional verbal versus non-verbal distinction. Instead, recent studies have focused on how these two modalities operate as components of a unified communicative system. From the 1980s onwards, gesture research has grown rapidly, particularly within the fields of psycholinguistics and cognitive psychology.

One key area of interest in gesture research is the function that gestures serve. Numerous studies have aimed to clarify these functions. In a review of earlier work, Kita (2000) identified two primary roles: conveying information and facilitating the processes of speech production and thinking. The former is known as the listener-oriented function, and the latter as the speaker-oriented function. The listener-oriented function refers to the role of gestures in conveying messages to listeners and supporting communication. For instance, studies have shown that listeners may reproduce information conveyed solely through a speaker’s gestures in their own speech (Cassell, McNeill, & McCullough, 1999).

Furthermore, even in the absence of formal training, teachers have been observed using children’s gestures as cues for assessment (Goldin-Meadow, Wein, & Chang, 1992). Observational studies have also suggested that gestures help maintain and regulate turn-taking in conversation. These findings provide converging evidence for the listener-oriented function of gestures (Bavelas et al., 1992).

In contrast, the speaker-oriented function of gestures concerns their role in facilitating the speaker’s own verbalisation and speech production. Evidence for this function includes findings that speakers continue to gesture even when speaking on the phone or when a physical barrier prevents visual contact with the listener (Alibali et al., 2001; Barroso, Freedman, & Grand, 1978; Rimé, 1982). Additionally, studies have shown that congenitally blind children and adults gesture during specific tasks in ways similar to sighted children and adults (Iverson & Goldin-Meadow, 1997; Mamus et al., 2023), suggesting that gestures serve a purpose beyond merely providing visual information to the listener. More recent empirical studies have focused on clarifying the mechanisms by which gestures support speech production. Research has demonstrated that gestures aid in concept formation (Alibali, Kita, & Young, 2000) and facilitate lexical retrieval (Krauss, Chen, & Gottesman, 2000; Rauscher, Krauss, & Chen, 1996), both of which offer further support for the speaker-oriented function of gestures.

Gesture–speech production models have been developed to explain how gestures are generated and how they relate to speech, thought, and communication processes. These models are primarily grounded in Levelt’s (1989) information-processing framework for speech production, which outlines a modular architecture (e.g., De Ruiter, 2000; Krauss et al., 2000). According to Levelt’s model, speech production involves a sequence of modular stages: the conceptualiser, formulator, and articulator. The conceptualiser generates ideas and passes them to the formulator. The output of the conceptualiser is a preverbal message, which is not yet in linguistic form but is instead represented in a non-verbal format. In the formulator, this preverbal message is transformed into a phonetic plan for articulation. This stage includes lexicalisation (assigning words to preverbal messages), grammatical encoding (structuring the message), and phonological encoding (spelling out a phonetic plan for the utterance). Finally, the articulator converts the phonetic plan into motor commands for the speech apparatus, such as the tongue, lips, and larynx. Proposed gesture-speech production models differ in terms of which stage of the speech production process gestures are thought to influence. For example, some models propose an effect at the stage of conceptualisation, while others emphasise lexical retrieval. Building on Levelt’s model, three major gesture-speech production models have been proposed, each presenting a hypothesis to explain how gestures may facilitate speech production.

The Image Activation Hypothesis proposes that gesturing enhances the activation of mental imagery associated with gestures, which in turn helps translate thoughts into verbal expressions (de Ruiter, 1998). This hypothesis assumes that gestures are connected to the conceptualiser in the speech production process. Empirical studies have shown that restricting bodily movements reduces the vividness of verbal descriptions, indicating a strong link between gestures and conceptual imagery (Rimé et al., 1984). Further findings support this view, demonstrating that limiting hand movements impairs the ability to describe concepts. This suggests that gesturing enhances a speaker’s mental imagery of the intended content and thereby facilitates more effective verbal formulation through image activation (Rimé, 1982).

An alternative explanation, known as the Lexical Retrieval Hypothesis, suggests that gestures support speech by aiding in word retrieval, particularly when speakers experience lexical access difficulties (Butterworth and Hadar, 1989; Hadar and Butterworth, 1997; Rauscher, Krauss, and Chen, 1996). This hypothesis assumes that gestures are linked to the formulator in the speech production process. Specifically, gestures are thought to convey concepts in a motoric or kinesic form to the phonological encoder within the formulator. This motoric representation facilitates access to word forms through cross-modal priming. Some studies have found that gesture use increases during pauses or moments of word-finding difficulty, as gestures often occur during hesitation pauses or immediately before problematic lexical retrieval (Dittmann and Llewelyn, 1969; Butterworth and Beattie, 1978; Theochaaropoulou et al., 2015). Other research has reported that restricting gesture use can result in verbal disfluency (Dobrogaev, 1929). These findings suggest that gesturing may enhance speech fluency by supporting lexical search. However, some scholars have challenged this view, arguing that there is no direct causal relationship between gesture inhibition and fluency in speech (e.g., Kısa et al., 2021; Graham and Heywood, 1975; Nobe, 2000; Hoetjes et al., 2014).

Lastly, the Information Packaging Hypothesis (Kita, 2000; Alibali et al., 2000) proposes that gestures are linked to the conceptualiser, where they assist in organising and structuring information prior to speech formulation. This hypothesis is supported by studies showing that speakers gesture more frequently when facing high cognitive demands during conceptualisation or when introducing novel content (Kita and Davies, 2009; Bergmann and Kopp, 2006). In addition, cross-linguistic evidence suggests that gestures reflect language-specific patterns of information structuring, further reinforcing the close link between gesture and speech (Kita and Özyürek, 2003).

While these gesture production models offer valuable insights into the role of gestures in speech production, none of them clearly defines the time course of gesture production in relation to speech. This limitation arises because previous studies have relied exclusively on behavioural indicators. Levelt’s speech production model, along with gesture production models based on this framework, suggests that speech generation operates primarily in a feedforward manner. In such a model, information flows largely in a single direction, from conceptualisation to articulation, with minimal dependence on immediate sensory feedback during the core stages of planning and execution. However, it remains unclear whether gestures influence online speech production before, during, or after articulation. If gestures truly facilitate conceptualisation or lexical retrieval in real time, as proposed by these models, they would need to be initiated well before the corresponding words. In practice, however, gestures and speech are typically produced simultaneously. In other words, they are temporally synchronised.

The synchrony between gesture and speech suggests that the speaker’s intention or mental preparation to gesture may be sufficient to support speech production. This raises an important and unresolved question: how can gestures influence speech before they are physically realised? To investigate this, we measured brain activity during gesture and speech production using magnetoencephalography (MEG). MEG allows for high temporal resolution recordings of brain activity, with a sampling rate of approximately 1000 Hz. This approach enabled us to examine the temporal dynamics of neural activity and identify the brain regions associated with both gesture use and speech production. To our knowledge, MEG has not previously been used to investigate gesture production, highlighting the academic originality of this study. By employing this method, we were able to provide the first evidence of real-time temporal information directly linked to the neural mechanisms involved in both gesture and speech production. Based on prior findings linking the anterior temporal lobes (ATLs) to lexical and semantic processing (e.g., Patterson et al., 2007; Lambon Ralph et al., 2017), we selected this region a priori to test whether gestures modulate pre-articulatory language planning.

Understanding the time course of gesture production is theoretically important precisely because existing accounts provide no explicit claims about the temporal dynamics of how gestures influence speech. Although the existing hypotheses differ in the cognitive mechanisms they prioritise, none specifies when, within the unfolding stages of speech production, gesture is expected to exert its influence. As a result, the models remain difficult to compare or empirically differentiate in their current form. Even though these accounts assume that gestures facilitate speech, gestures typically emerge at almost exactly the same moment as the corresponding speech. Thus, any facilitative effect must arise earlier, during the preparation or planning stages of gesture production, rather than at the moment the gesture becomes visible. Such temporal information cannot be obtained from behavioural data alone. By leveraging MEG’s millisecond-level resolution, the present study provides the first opportunity to clarify the timing of gesture–speech interactions and, in doing so, to render the existing theoretical accounts more precise and empirically distinguishable.

Previous research on the neural basis of gesture has focused primarily on gesture and speech comprehension. Studies have examined whether listeners or observers can extract meaning from gestures and speech, whether they integrate these two modalities, and which brain regions are involved in processing them (e.g., Demir-Lira et al., 2018). Some research has investigated the online processing of gesture and speech using electrophysiological measures, such as event-related potentials (ERPs) (e.g., Drijvers & Özyürek, 2018; Habets et al., 2011; Holle and Gunter, 2007; Sekine et al., 2020).

ERPs are measured using electroencephalography (EEG) and provide precise temporal information about the timing of neural processing. Research on gesture-speech integration often focuses on the N400 effect, which is known to reflect the ease of semantic integration between a word and its preceding context, or between a word and a gesture (Drijvers & Özyürek, 2018). It is also associated with activation in left-lateralised front-temporal regions (Özyürek, 2014). The N400 effect refers to a negative-going deflection in the ERP waveform occurring between 300 and 550 milliseconds after stimulus onset, with increased amplitude in response to semantically incongruent information in words or gestures, compared to congruent ones. For example, using this measure, Kelly et al. (2004) examined the neural correlates of speech and iconic gesture comprehension. They found that ERPs elicited by spoken words (targets) were modulated when these words were accompanied by gestures (primes) that conveyed information about the size or shape of the referent objects (e.g. tall, wide). Compared to matching targets, mismatching words evoked a stronger N400 response, suggesting that gestures influence spoken word processing at the semantic level. Similarly, Habets et al. (2011) examined whether the temporal alignment of gesture and speech affects integration. In their study, video stimuli paired gestures with either congruent (e.g. connecting) or incongruent (e.g. falling) verbs, presented at three levels of synchrony: 0ms (fully synchronous), 160ms (partially overlapping), and 360ms (not overlapping). ERPs time-locked to speech revealed significant N400 differences between congruent and incongruent conditions at 0ms and 160ms, but not at 360ms. These results suggest that the closer the temporal alignment between speech and gesture, the more likely they are to be semantically integrated.

In contrast to EEG studies, which reveal when the integration process occurs, functional magnetic resonance imaging (fMRI) identifies where the process takes place. Using fMRI, several studies have attempted to locate the brain regions in adults that are involved in integrating information from speech and gesture. To identify these regions, researchers typically compare multimodal stimuli with unimodal ones, or gestures presented with degraded speech with those presented alongside clear speech. Findings suggest that the integration of iconic gestures and speech in adults recruits brain regions such as the left inferior frontal gyrus (IFG), posterior superior temporal sulcus (STSp), middle temporal gyrus (MTGp), and even the motor cortex (e.g. Holle et al., 2008; Özyürek et al., 2007; Straube et al., 2009; Willems et al., 2008). These areas are also known to be involved in the semantic processing of speech comprehension (Dick et al., 2008, 2012). In particular, studies have highlighted the importance of posterior temporal regions, especially STSp and MTGp, in the semantic integration of gesture and speech. Green et al. (2009) found that the left MTGp showed stronger responses to sentences paired with unrelated gestures, which were difficult to interpret in relation to the speech content, than to the same sentences accompanied by related gestures. Similarly, Holle et al. (2008) reported that STSp was more active when ambiguous words were accompanied by meaningful iconic gestures than when paired with non-meaningful grooming movements. Additionally, Holle et al. (2010) found that STSp showed greater activation when gestures were presented with degraded speech, compared to clear speech.

These findings suggest that the STSp plays a key role in the semantic integration of gesture and speech. Moreover, STSp is known to be sensitive not only to gesture-speech integration but also to other forms of multimodal integration, such as the combination of lip movements and syllables (Willems et al., 2008). However, the nature of the online processes underlying the production of gestures and speech remains unclear. To date, only a few studies have investigated gesture production using neuroimaging or neurophysiological techniques (e.g. Oi et al., 2013; Sato et al., 2023; Wang et al., 2024).

Oi et al. (2013) used near-infrared spectroscopy to investigate spontaneous co-speech gesture production during an animation narration task in bilingual speakers. They reported increased gesture use and greater activation in the left inferior frontal gyrus under higher cognitive load, particularly when participants spoke in their second language, consistent with a facilitative role of gesture in speech production. While this study provided important neural evidence for gesture-related support during language production and informed aspects of the present experimental design, it did not address the temporal dynamics of how gestures influence speech planning. More recently, Wang et al. (2024) employed fNIRS hyperscanning to examine spontaneous gestural communication in interacting dyads, showing enhanced inter-brain network efficiency when gestures were allowed. Neural alignment was strongest within mirroring and mentalising networks, including regions such as the inferior frontal gyrus, superior temporal gyrus, temporo-parietal junction, and prefrontal cortex, highlighting a focus on interpersonal coordination rather than on the timing of gesture effects on individual speech planning. Together, these findings emphasise the role of gestures in interpersonal neural coordination, rather than in the temporal dynamics of individual speech planning that are the focus of the present study.

Lastly, Sato et al. (2023) investigated the neural basis of gesture and speech planning using EEG effective connectivity analysis. Participants were asked to plan either a gesture or a spoken word in response to spatial images while their brain activity was recorded. The results showed that gesture planning elicited stronger connectivity from the right middle cingulate gyrus (MCG) to the left supplementary motor area (SMA), and from the left SMA to the left precentral area (PCA), compared to speech planning. These findings suggest that gesture production relies more heavily on motor-related neural networks, distinguishing its neural dynamics from those involved in speech planning.

All three gesture production studies support previous fMRI findings on gesture comprehension, indicating that the left IFG is crucial not only for understanding gestures but also for producing them. These results reinforce the link between gesture and speech in language processing. However, none of these gesture production studies has explored the temporal dynamics between speech and co-speech gestures. Instead, they have focused either on identifying the brain areas involved in gesture production or on brain activity during gesture planning without actual execution. Therefore, the current study is the first to investigate the time course of gesture production, specifically in terms of when gestures influence speech.

In the present study, we specifically examined whether allowing co-speech gestures modulates neural activity during the planning phase of speech production. To this end, participants performed a spontaneous narration task under two conditions: a Gesture-Required condition and a Gesture-Prohibited condition. Using speech onset as a temporal reference point, we tested whether gesture-related neural modulation emerges prior to articulation. Based on prior evidence implicating the anterior temporal lobes in lexical-semantic processing, we hypothesised that permitting gestures would attenuate neural activity in these regions during the pre-articulatory period. In contrast, if gesture-related effects primarily reflect motor preparation, comparable modulation would be expected in motor-related regions.

## 2. Method

### 2.1 Participants

Twenty-six participants (all native Japanese speakers, three females, mean age = 23.2 years, SD = 1.8) took part in this study. Three participants (all males) were excluded from the subsequent analyses either because they were unable to complete the required number of trials due to MEG setup issues, or because some of their vocal responses were not properly recorded. These exclusion criteria were established prior to the analysis of any MEG data. The group size was determined based on a power analysis using data from one of the authors’ previous MEG studies (Chang et al., 2021). All participants had normal or corrected-to-normal vision and no history of neurological disorders. They gave both written and oral informed consent prior to MEG recording. All experimental procedures were approved in advance by the ethics committee of the National Institute of Information and Communications Technology (NICT), Japan (N23006240), and the CiNet safety committee (2405150120).

### 2.2. Experiment design and tasks

During the MEG acquisitions, participants performed a task that involved watching short silent video clips (1280 × 720 pixels, 29.97 frames per second each) and describing their contents verbally, accompanied by gestures, one at a time (Figure 1). The task was controlled using MATLAB Psychtoolbox (http://psychtoolbox.org/) and custom in-house subroutines, running on a PC (QX3P5000 with an NVIDIA Quadro P5000 graphics card, Zotac, Hong Kong, China). The video clips were back-projected via an LCD projector (WUX400, Canon, Japan) onto a translucent screen (movie clip size: 31.4 × 23.6 degrees of visual angle) placed in front of the participant’s forehead.

**Figure 1.**
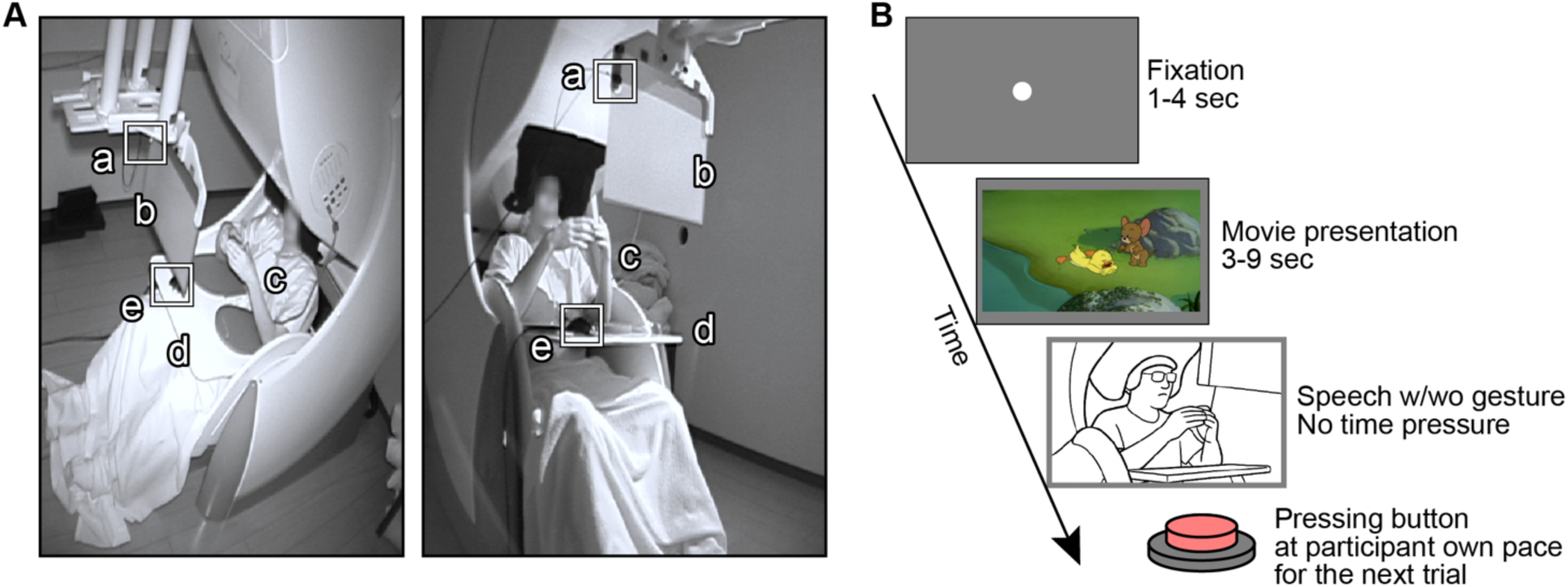
MEG measurements. (A) MEG recording setup. Entire MEG sessions were audiovisually recorded using an infrared camera. (a) Photodiode used to mark video onset and offset, (b) Visual stimuli were presented on this screen, (c) Gestures were performed mildly using only the forearms and hands, (d) A wooden board was mounted to the MEG enclosure and placed between the participant’s lap and arms. Participants were instructed to keep their elbows on this board, (e) Response box used to initiate the next trial. (B) Example of a trial sequence.

In each trial, participants viewed a 3- to 9-second silent film clip composed of various segments from the Tom and Jerry cartoon series (Warner Bros. Entertainment, Inc., USA). Participants were instructed to imagine sitting next to a friend and to briefly explain the content of the clip they had just watched, within approximately 10 seconds. They were also asked to use gestures along with verbal descriptions for the Gesture-Required condition. Participants were instructed to narrate the content of the cartoon in an everyday, conversational manner, and no explicit instructions were given regarding the frequency with which gestures should be produced across utterances. In this condition, to minimise motion artefacts in the MEG signal caused by hand movements, participants were instructed to keep their elbows on a wooden board and to perform only mild gestures involving the forearms, wrists, and hands, without lifting their elbows from the board (Figure 1A). After completing each explanation, participants pressed a button to initiate the next trial at their own pace.

In the Gesture-Prohibited condition, which served as a control, participants were instructed to verbally describe the video content without using any gestures, while keeping their hands still and resting on a wooden board mounted to the MEG enclosure and placed between their laps and arms. To avoid confusion and to prevent movement-related artefacts caused by repeated hand placement and gesture transitions, the two task conditions were conducted in separate MEG measurement sessions, with no intermixing within a single session. Each session consisted of 30 trials of one task condition and lasted approximately 12 minutes on average across participants. For each condition, participants completed three or four separate recording sessions, yielding a total of 90 to 120 trials per condition. The order of the stimulus conditions was counterbalanced across participants.

### 2.3. MEG acquisition

MEG data were recorded in a magnetically shielded room at the NICT CiNet imaging facility (Suita, Osaka, Japan), using a customised 360-channel whole-head MEG system (Elekta Neuromag 360, MEGIN Oy, Helsinki, Finland) (see Figure 1). The system comprised 204 planar gradiometers, 102 magnetometers, and 54 additional sensors (18 magnetometers and 36 gradiometers, all vertically placed to detect tangential magnetic field components).

Magnetic signals were originally sampled at 5000 Hz and were downsampled to 1000 Hz during recording. A participant’s head position relative to the sensor array was continuously tracked using five head-position indicator coils attached to the forehead and preauricular points. The precise locations of these coils were digitised prior to recording using a 3D digitiser.

The timing of MEG signal acquisition and video presentation was synchronised offline on a trial-by-trial basis using a photosensor (MRI-OE2-ATR, Newopto, Kanagawa, Japan). The sensor was positioned at the upper-right corner of the screen and detected the onset of a white square probe presented during the movie presentation periods. To prevent participants from seeing the synchronisation marker, the probe and the photosensor were

All MEG measurement sessions were audiovisually recorded using an infrared video camera, with prior informed consent obtained from all participants. These recordings were used to annotate the onsets of verbal responses and gestures offline using ELAN software (Lausberg & Sloetjes, 2009). To enable precise temporal alignment between behavioural annotations and MEG data, an audio marker consisting of three successive high-tone beeps was introduced at the beginning of each MEG recording session. This audio signal was converted into an electrical signal and recorded via an external MEG channel, and was also captured by the video recording. Offline alignment was performed by matching the audio waveform recorded by the MEG system with the corresponding audio track in the video recordings, thereby minimising temporal discrepancies between annotated events and the MEG signal.

### 2.4. Behavioural data (video) annotations and analysis

As described above, the MEG recording sessions were audiovisually recorded using an infrared camera, and behavioural events were manually annotated offline based on the video footage. Through this process, the precise timings of the following events were identified: the onsets of the triple high-pitched beeps, the onsets and offsets of the Tom and Jerry video clips, the onsets and offsets of gestures, and the onsets and offsets of verbal responses. These annotated timestamps were subsequently used in the behavioural analysis to examine timing and duration differences between the Gesture-Required and Gesture-Prohibited conditions.

### 2.5. MEG data preprocessing

The raw MEG signals were primarily pre-processed using the following three steps. First, Oversampled Temporal Projection (OTP) (Larson & Taulu, 2018) was applied to attenuate temporally uncorrelated high-frequency noise components in the time domain. Second, the XSCAN tool (MEGIN Oy, Helsinki, Finland) was used to identify and flag channel-specific artefacts, determining which channels should be excluded during subsequent signal reconstruction as “bad channels.” Finally, the temporal extension of Signal Space Separation (tSSS), implemented via the MaxFilter software (MEGIN Oy, Helsinki, Finland), was applied to suppress external noise sources (Taulu et al., 2005; Taulu & Hari, 2009). The tSSS algorithm achieves this by temporally and spatially separating magnetic field components originating from inside and outside the sensor array, using an expansion of the signal vectors into spherical harmonics. For automatic artefact rejection, a buffer length of 10 seconds was employed, following the default parameters described in the official tSSS documentation.

The pre-processed MEG signals were notch-filtered at 60, 120, 240, 360, and 420 Hz to remove power-line noise. They were then downsampled to 200 Hz and band-pass filtered between 0.1 and 45 Hz. Next, the MEG signals underwent an ICA-based decomposition (Hyvärinen, 1999), and artefactual components derived from ocular activity (e.g. blinks and saccades) and cardiac activity (e.g. breathing and heartbeats) were automatically identified and removed during signal reconstruction. Finally, the MEG time series were normalised by subtracting the global mean of the signal for each channel within each recording session.

### 2.6. MEG data analyses

All subsequent analyses were conducted using the MNE-Python software package (Gramfort et al., 2013; https://www.martinos.org/mne/stable/index.html) along with custom in-house subroutines. For the sensor-level ROI analyses, only MEG signals recorded by the magnetometers were used unless otherwise specified (see Figure S2 in the supplementary materials). First, to obtain an overview of the MEG response profiles aligned to the onset and offset of verbal responses or gestures, we aggregated data from all channels and computed root mean square (RMS) responses. This analysis also served as a sanity check. In conventional MEG recordings, participants are typically required to remain still, as any movement of the hands, feet, or head can introduce substantial noise or artefacts into the data. However, because the aim of the present study was to investigate how gestures influence speech production, it was necessary to permit hand movements during MEG acquisition. Based on pilot testing, we found that asking participants to keep their elbows on a wooden plate mounted between their laps and arms, and to perform only mild gestures using the forearms and hands without lifting the elbows, helped preserve the naturalness of the gestures while minimising motion-related artefacts.

The results of the RMS analysis confirmed that MEG signal changes were reliably observed around the onset of speech. This indicated that, despite the potential risks associated with allowing gestures during MEG acquisition, our procedure, which constrained gestures to a limited and controlled range, successfully mitigated concerns related to motion artefacts. Crucially, the RMS analysis aligned to speech onset revealed that gesture-related neural effects emerged prior to the initiation of speech. Based on this finding, subsequent analyses were conducted at the individual sensor level to investigate both the magnitude of responses and the spatial distribution of activity during the pre- and post-speech periods. Based on prior literature (e.g., Oi et al., 2013), bilateral anterior temporal lobe and motor cortex sensor regions of interest were defined a priori; subsequent condition-wise comparisons of neural responses were conducted exclusively within these predefined ROIs.

## 3. Results

### 3.1 Behavioural Results: Gesture enhances verbal elaboration

We first examined whether the use of gestures leads to more elaborate speech production at the behavioural level. If gestures support conceptualisation or lexical retrieval, as suggested by gesture production models based on Levelt’s framework, participants should produce longer or more detailed utterances when gestures are permitted. We thus compared the speech durations between the two experimental conditions. For each participant, we calculated the durations from speech onset to offset in all the trials and then averaged these durations separately for the Gesture-Required and Gesture-Prohibited conditions. A paired-samples t-test confirmed that mean speech duration was significantly greater when gestures were allowed (M = 9.03 seconds, SD = 3.03) than when gestures were prohibited (M = 7.52 seconds, SD = 2.68), *t*(22) = 3.43, *p* = .002. This finding indicates that the availability of hand gestures during narrations led to more elaborated verbal descriptions, suggesting that gestures may facilitate the planning or release of additional linguistic content.

Additional behavioural analysis revealed that gesture onsets were approximately normally distributed (based on visual inspection) around the onset of speech, with a mean lag of approximately zero delay (Supplementary Figure S1). This finding indicates that gestures and speech were, on average, initiated simultaneously. The temporal alignment between gesture and speech supports the view that these two modalities are co-planned rather than independently initiated. Importantly, it also suggests that the MEG signal differences observed in the 250 ms preceding speech onset (the details are described in the later section) are unlikely to reflect preparatory motor activity specific to gestural execution as it may be generally too early as a motor preparation activity. Instead, they are more plausibly attributed to pre-articulatory cognitive language processes such as lexical retrieval or information packaging.

However, due to methodological constraints specific to MEG acquisition, we were unable to include additional linguistic measures such as word counts or syntactic complexity. The only microphone available at the time that did not interfere with the MEG sensors produced low-fidelity audio. Although the recordings were sufficient for detecting speech onsets and offsets, the audio quality was inadequate for reliably identifying individual words or transcribing utterances. For this reason, speech duration was used as the sole behavioural index in the present study.

### 3.2 MEG Results: Gesture-derived MEG signal attenuation precedes speech onset

The behavioural results raised the question of which neural mechanisms underlie this speech facilitation and how gestures modulate cortical activity. To answer these questions, we analysed the obtained high-temporal resolution MEG brain response patterns (for the details of the sensor arrays, please see Supplementary Figure S2) in the following two steps. Firstly, we visualised and compared the magnetic field topographic maps between the two conditions to identify when the gesture effect can be observed as the whole brain response pattern differences. Second, we computed the grand-averaged root mean square (RMS) signals across all magnetometer channels, time-locked to speech onset (from –2 to +1 seconds relative to speech onset at 0.1-second intervals), and explored when relative to speech onset the gesture effect is observed.

Visual inspection of the topographic maps (Figure 3) revealed a divergence in activity between the Gesture-Required and Gesture-Prohibited conditions shortly before speech articulation, approximately between −0.3 and +0.1 s relative to speech onset. This preliminary visualization guided the selection of a narrower time window for the latter quantitative analysis.

**Figure 2.**
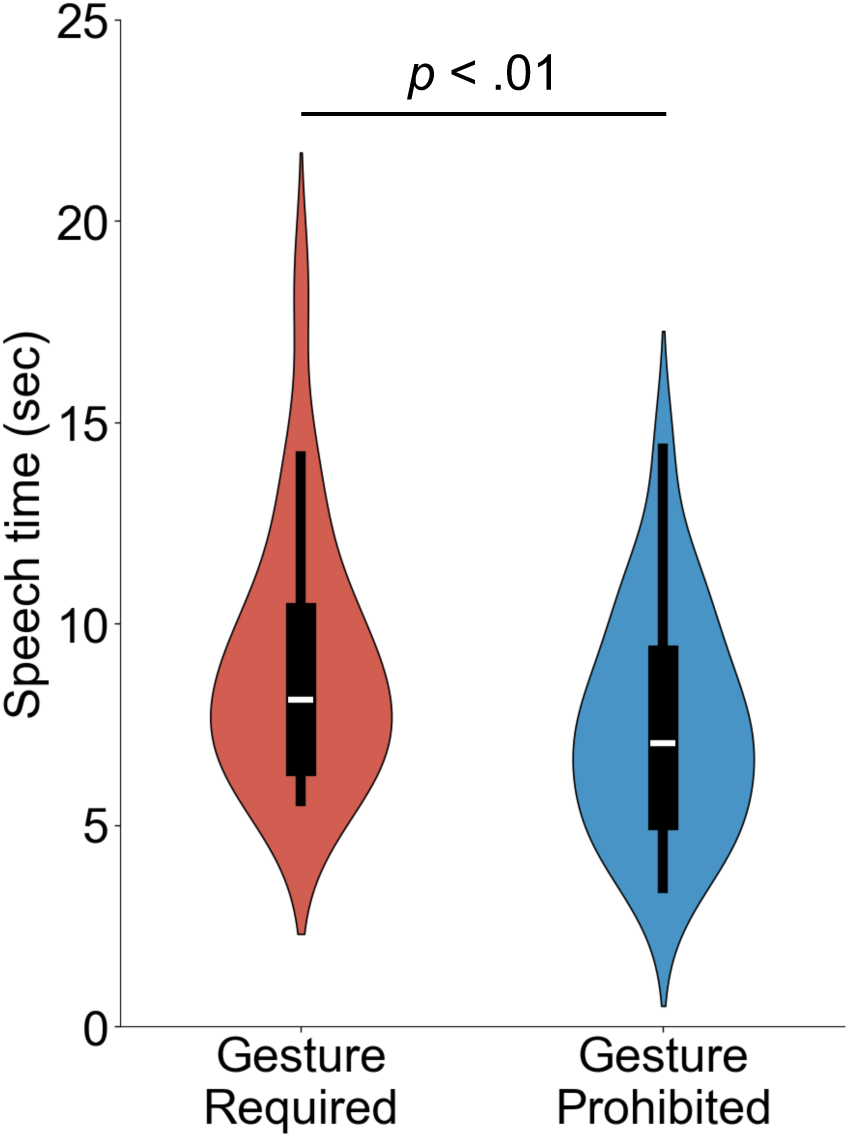
Speech time under the two conditions: Speech duration was significantly longer in the Gesture-Required condition than in the Gesture-Prohibited condition

**Figure 3.**
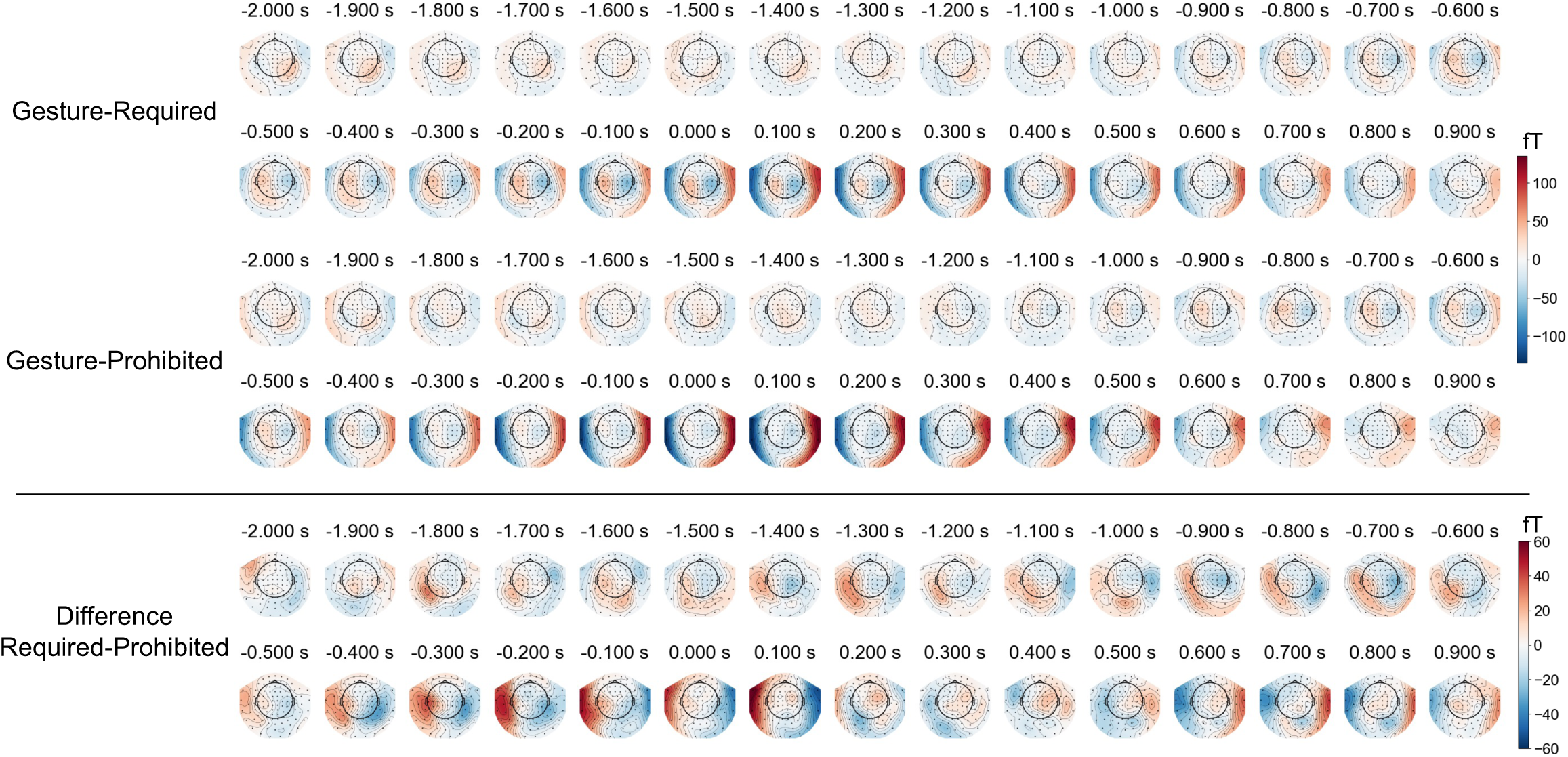
Time series of topographic maps around speech onset (−2 to 1 seconds). Time 0 indicates speech onset. A difference between the conditions was observed from −0.25 to 0 seconds. Top: Gesture-Required condition; Middle: Gesture-Prohibited condition; Bottom: Difference between the two conditions. Due to the three-dimensional, helmet-shaped structure of the MEG device, some channels may appear to lie outside the head when projected onto a two-dimensional plane.

The RMS signal comparisons showed that the response divergence was observed preceding the speech onset (Figure 4). Specifically, we computed RMSs of magnetometer signals across all channels for each condition and aligned them to speech onset. The RMS time course exhibited a clear attenuation for the Gesture-Required compared to the Gesture-Prohibited conditions in the –0.25 to 0 second window. This analysis served both as a sanity check for data quality and as a basis for identifying the pre-speech period as the temporal window of interest (Oi et al., 2013), and as a basis for defining a temporal window of interest (from -0.25 to 0 seconds) for subsequent statistical comparisons.

**Figure 4.**
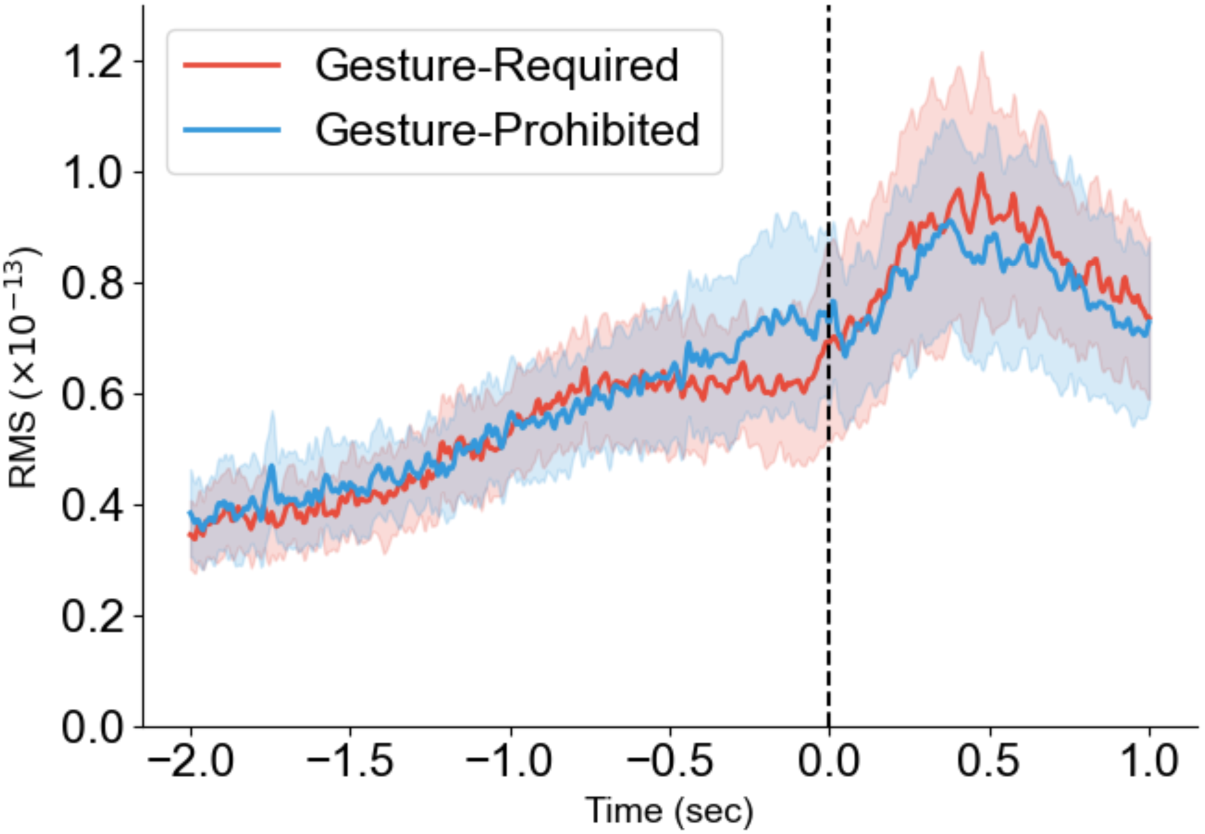
Root mean square (RMS) aligned to speech onset. A difference between the Gesture-Required and Gesture-Prohibited conditions was observed between −0.25 and 0 seconds. No statistical tests were conducted. The colored fill represents the 95% confidence interval.

We further showed that gesture-related response attenuation was localised in the anterior temporal lobes (ATLs). In this analysis, we conducted sensor-based response comparison during the predefined temporal window of interest. Based on prior evidence implicating the anterior temporal lobes (ATLs) in semantic and lexical processing, we hypothesised that the gesture effect would be detected over sensor regions corresponding to the bilateral ATLs as regions of interest (ROIs). Practically, we selected four magnetometer sensors located anatomically close to the right or left ATL region as shown in Figure 5A and contrasted mean signal amplitudes between conditions during the identified -0.25 to 0 s window. Motor-related regions (four magnetometer sensors close to the right or left anterior regions shown in Figure 5A) were also included as a control ROI to rule out non-linguistic sources of variation, such as hand movements associated with gestures and mouth movements associated with speech.

**Figure 5.**
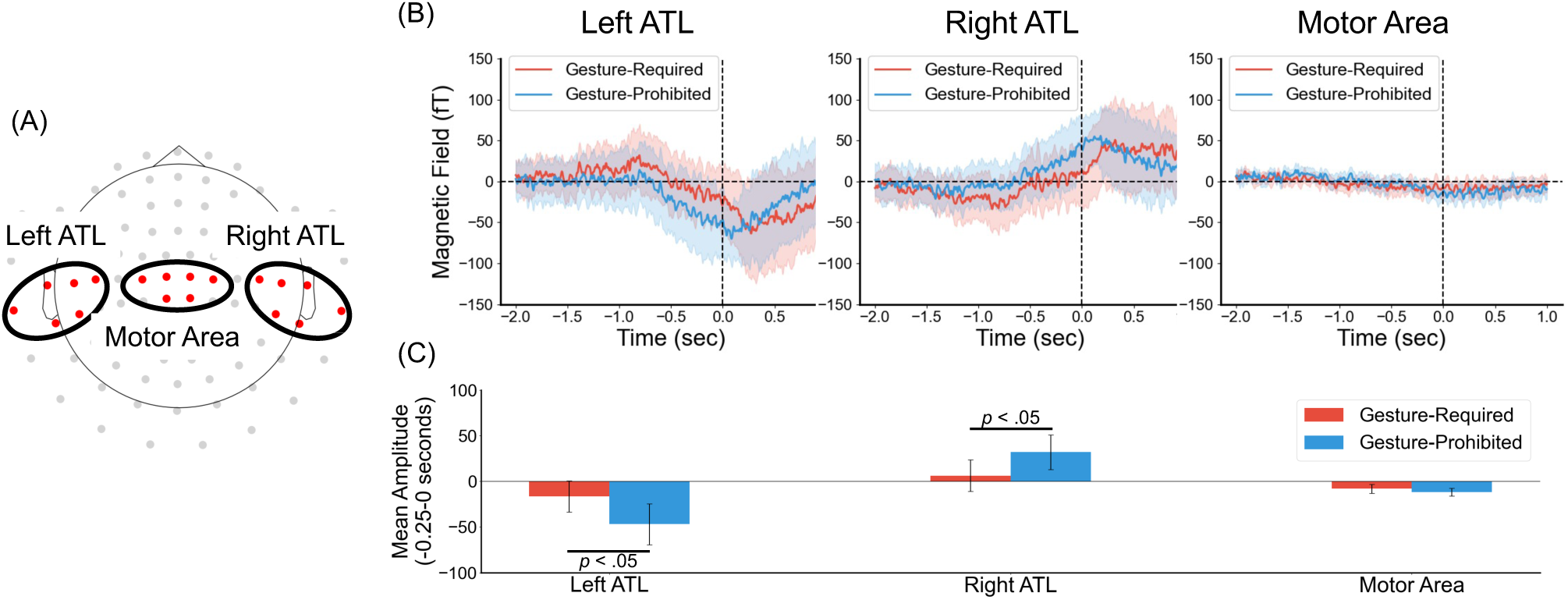
Event-related fields (ERFs) by region of interest (ROI). (A) Channels defined for each ROI. (B) Average waveforms for each ROI: left ATL (left), right ATL (centre), and motor area (right). (C) Mean amplitude between −0.25 and 0 seconds for each ROI. Time 0 indicates speech onset. In both the left and right ATL, the amplitude between −0.25 and 0 seconds was significantly lower in the Gesture-Required condition than in the Gesture-Prohibited condition. No significant differences were observed in the motor area. The shaded areas in panel B represent 95% confidence intervals, and the error bars in panel C indicate standard deviations.

It is worth clarifying here that, while the results are derived from the same dataset, the ROI (sensor)-based temporal-window-of-interest comparisons were not circular or post hoc in nature. The RMS analysis identified a time window (−0.25 to 0 s) showing condition differences in overall signal magnitude; however, it did not inform us about the spatial origins of this effect. The subsequent ROI-based temporal-window-of-interest analysis was conducted to determine where in the brain these differences arose, based on a priori hypotheses informed by prior literature implicating the anterior temporal lobes (ATLs) in semantic and lexical processes. Therefore, the RMS and ROI-based analyses thus address distinct but complementary questions, namely, temporal localization of gesture effect versus spatial identification of their cortical sources and together provide converging evidence for gesture-related modulation of pre-speech neural activity.

As shown in Figure 5(B) and 5(C), event-related responses were observed in both ATLs around speech onset. Difference between the conditions were also observed. Paired-samples t-tests revealed significantly greater ATL activity in the Gesture-Prohibited condition compared to the Gesture-Required condition (Left ATL: *t*(22) = 2.08, *p* = .049; Right ATL: *t*(22) = 2.35, *p* = .028). These findings support the hypothesis that gestures reduce cognitive load in language-related regions during the preparatory phase of speech production.

An alternative possibility is that the response attenuation in ATLs during the pre-speech period may be due to some simple motor preparation effect. This interpretation is plausible because speech and gesture occurred at approximately the same time, as shown in Supplementary Figure S1. If the ATL attenuation simply reflected preparation for hand movement, a motor-preparation effect should have been observed before speech onset in the Gesture-Required condition. However, the motor area did not show any clear response patterns as shown in Figure 5B. Furthermore, no significant conditional difference was observed in the motor ROI (Figure 5C, *t*(22) = 1.08, *p* = .291). These results make a simple motor artefact account unlikely.

To ensure that the observed effects were not widespread across the cortex outside the predefined ATLs, we conducted a complementary sensor-level analysis surveying the full sensor array (Supplementary Figure S3). This analysis confirmed that significant gesture-related differences were confined to the bilateral ATL regions, with no comparable effects observed in other sensor locations, including frontal, parietal, or occipital areas. Together with the absence of modulation in the motor control ROI, this supports the spatial specificity of gesture-related neural modulation within language-relevant temporal regions.

As a control analysis, we also examined MEG signals aligned to the onset of video presentation rather than speech (Supplementary Figure S4 and S5). In this alignment, no reliable differences between conditions were observed. This absence of effect suggests that the timing of gesture- and speech-related neural activity was not locked to the external stimulus (i.e., video onset), but rather to internally generated events such as speech initiation. This finding reinforces the interpretation that gesture-related neural modulation occurs during speech planning, not during visual processing or passive stimulus viewing.

To clarify the functional nature of the observed effects, we investigated whether the gesture-induced modulation reflected linguistic planning rather than general motor preparation. The absence of significant differences in motor cortex activity, alongside robust effects in the anterior temporal lobes (ATLs), supports a domain-specific interpretation: co-speech gestures facilitate conceptual or lexical processing without triggering preparatory motor signals unrelated to speech. These findings reinforce the view that gestures are not merely motor accompaniments but serve as integral components of the speech planning system, operating at a linguistic and semantic level. In sum, our results demonstrate that gesture use is associated with reduced neural activity in language-related cortical regions during the preparatory phase of speech, supporting the notion that gestures fulfil a cognitive-linguistic, rather than purely motor, function in spontaneous language production.

## 4. Discussion

Our findings reveal that co-speech gestures modulate neural activity in speech-related brain regions even before the onset of speech, specifically in the bilateral anterior temporal lobes (ATLs). Notably, activity in these regions was significantly reduced in the Gesture-Required condition compared to the Gesture-Prohibited condition during the 250ms window preceding speech onset. No significant differences were observed in motor areas, suggesting that these effects are unlikely to be driven by general motor preparation. Although previous studies have highlighted regions such as the IFG, STS, and MTG in gesture–speech integration, these findings mainly reflect comprehension processes (e.g. Holle et al., 2008; Özyürek et al., 2007; Willems et al., 2008). In contrast, our study focused on gesture effects during speech production, particularly in the planning phase. Based on their established role in semantic and lexical access, we targeted the anterior temporal lobes (ATLs) as the primary ROI.

Together, these results provide direct neurophysiological evidence that gestures play a facilitative role during the speech planning phase, thereby supporting speaker-oriented models of gesture production (e.g., Kita, 2000; Alibali et al., 2000; Krauss et al., 2000). In particular, the observed reduction in ATL activity aligns with the Lexical Retrieval Hypothesis (Rauscher et al., 1996), which posits that gestures assist in retrieving lexical items by activating semantic representations in a motoric or imagistic format. Reduced ATL activity in the Gesture-Required condition may reflect decreased processing demands during lexical access, potentially because gestures had already primed relevant semantic fields (e.g., Alibali et al., 2000; Rauscher et al., 1996; Krauss & Hadar, 1999).

The anterior temporal lobes are widely regarded as a semantic hub supporting lexical and conceptual processing. Neuropsychological and neuroimaging evidence suggests that the ATLs support semantic memory and concept-level processing and are particularly involved in retrieving word meanings and resolving semantic ambiguity during speech production (Patterson et al., 2007; Lambon Ralph et al., 2017). In the present study, reduced ATL activity in the Gesture-Required condition likely reflects a decrease in semantic processing demands, suggesting that gestures may aid in semantic access or conceptual structuring before articulation.

Our findings are also consistent with the Information Packaging Hypothesis (Kita, 2000), which argues that gestures help speakers organise and structure complex information during conceptual planning. The anterior temporal lobes have previously been implicated in high-level semantic integration and conceptual combination, and reduced activity in these regions may indicate that gestures serve to offload conceptual structuring demands prior to speech formulation. Additionally, the results offer an extension of the Image Activation Hypothesis (de Ruiter, 1998), which emphasises the role of gesture in activating mental imagery. While earlier behavioural studies have inferred this function from speech content or fluency, our MEG data provide direct neurophysiological evidence that gestures influence the speech production process even before articulation, possibly by enhancing conceptual availability or activating visuospatial representations.

An important question raised by the present findings is whether they help adjudicate between existing theories of gesture production. Previous behavioural studies have suggested that gestures support speech planning, but have been unable to determine when this influence arises due to the near-synchronous timing of gesture and speech (McNeill, 1992; Nobe, 2000). Importantly, the present study was not designed to directly contrast these accounts.

Instead, it addresses a theoretical dimension that has remained largely underspecified across major models, namely when gestures influence speech production. Although the Image Activation, Lexical Retrieval, and Information Packaging hypotheses differ in the mechanisms they emphasise, they all assume that gestures facilitate speech planning, without specifying the timing of this influence. By showing reduced neural activity in the bilateral anterior temporal lobes during the 250 ms preceding articulation, the present study provides the first neurophysiological constraint on this shared assumption. The key theoretical contribution therefore lies not in selecting a single account, but in demonstrating that any viable model must allow for a pre-articulatory influence of gesture on language planning.

Importantly, the present analyses were conducted at the level of entire verbal responses rather than individual gesture–lexical pairings. As such, the observed effects should be interpreted as reflecting facilitation of global lexical-semantic planning or information packaging, rather than item-specific lexical retrieval. While the present results do not rule out imagery-based processes, they indicate that such contributions must operate prior to overt articulation rather than at the level of gesture execution. By specifying when gestures influence speech, this study refines existing models and renders them more precise and empirically testable.

Crucially, the absence of significant differences in motor regions suggests that the effects of gesture are not attributable to general motor preparation or execution but are instead specific to speech-related processing. This dissociation supports the view that the primary function of gesture in speech production is not merely biomechanical or expressive, but rather cognitive and linguistic in nature. Sato et al. (2023) found that the left supplementary motor area was more activated during gesture planning than during word planning. However, unlike our study, which focused on co-speech gestures, Sato et al. instructed participants to produce either a gesture or a word, but not both. Therefore, participants in their study likely paid more attention to hand movements in the gesture condition to represent a referent, which may have led to greater activation in the motor area compared to the speech condition.

Prior neuroimaging studies of gesture production have primarily focused on spatial localisation using fMRI or fNIRS, or have examined gesture comprehension rather than production (e.g. Oi et al., 2013; Sato et al., 2023; Wang et al., 2024). By leveraging the temporal precision of MEG, we captured the real-time dynamics of gesture–speech interaction and demonstrated that gestures modulate preparatory neural activity ahead of speech. This provides new insights into the timing of gesture’s influence on speech production.

While our findings contribute to understanding the predictive mechanisms involved in speech production, several limitations of the present study should be acknowledged. First, speech duration was used as the sole behavioural index of verbal output. This measure is relatively coarse and does not directly capture finer-grained linguistic properties such as lexical diversity, syntactic complexity, or discourse structure. Accordingly, the present findings should not be interpreted as evidence that gestures necessarily enhance linguistic elaboration in a narrow sense. Rather, longer speech durations in the Gesture-Required condition are more plausibly interpreted as reflecting reduced planning demands or greater ease of message formulation when gestures are available.

Second, although the Gesture-Required and Gesture-Prohibited conditions were counterbalanced, it remains possible that the explicit instruction to gesture or refrain from gesturing introduced unnatural cognitive strategies. Future work should therefore include more ecologically valid control conditions in which no explicit constraints on hand movement are imposed. Third, while the anterior temporal lobes emerged as a key region of interest, further source localisation techniques such as beamforming would allow for more precise spatial characterisation of the underlying neural generators. Fourth, although the present study involved spontaneous storytelling in Japanese, future research should examine whether these findings generalise across different languages and discourse contexts, particularly those with diverse typological features.

Finally, the present study could not examine whether the temporal relationship between gesture onset and speech onset varied as a function of gesture type such as iconic, metaphorical or beat gestures (McNeill, 1992). Although speech onset timing was available, reliably classifying gesture types requires identifying the specific spoken content with which gestures are synchronised. In the present dataset, this was not always possible due to variability in recording quality, which in some cases made the spoken content difficult to discern. Future studies combining tasks that elicit a broader range of gesture types with consistently high-quality audio recordings will be necessary to directly test whether different gesture types exert distinct pre-articulatory effects on speech planning. Importantly, existing theories that link gesture production to lexical and semantic facilitation are primarily grounded in gestures that carry representational content, such as iconic and deictic gestures. Consequently, the present findings should be interpreted with caution, as the observed facilitative effects may not generalise to gesture types with primarily pragmatic or interactive functions, which are not theoretically expected to engage semantic or lexical processes to the same extent.

In sum, this study provides the first MEG-based evidence that gestures reduce neural processing load in speech-related brain regions prior to speech, reinforcing the idea that gestures function as an integral component of the speech production system. Our findings support models that conceptualise gesture and speech as a unified, temporally coordinated system, and pave the way for further investigations into the neural mechanisms by which gesture scaffolds speech production.

## Acknowledgments

This study was supported by a research grant from the Waseda University–NICT Matching Research Support Program for the first and third authors, and by the Japan Society for the Promotion of Science (KAKENHI Grant Number 24K03239) for the first author. We thank N. Murakami and M. Hayashi for their support in organising the MEG experiments and data acquisition, and H. Nishimoto, A. Otsuka, and H. Mimura for conducting the MEG measurements and maintenance.

## Author approvals

All authors have seen and approved the manuscript. The manuscript has not been accepted for publication or published elsewhere.

## Author contributions

K.S. conceptualised the study and designed the experiment. R.O. and H.B. contributed to data acquisition and interpretation of the results. R.O. and H.B. contributed to MEG recording, data processing, and data analysis. K.S. wrote the original draft. All authors reviewed and edited the manuscript and approved the final version.

## Data availability

The datasets generated for the current study are available from the corresponding author upon reasonable request.

## Ethics Statement

This study was approved by the ethics committee of the National Institute of Information and Communications Technology (NICT), Japan (N23006240), and the CiNet safety committee (2405150120).

## References

Alibali, M. W., Heath, D. C., & Myers, H. J. (2001). Effects of visibility between speaker and listener on gesture production: Some gestures are meant to be seen. Journal of Memory and Language, 44(2), 169–188.

Alibali, M. W., Kita, S., & Young, A. J. (2000). Gesture and the process of speech production: We think, therefore we gesture. Language and Cognitive Processes, 15(6), 593–613.

Barroso, F. X., Freedman, N., & Grand, S. (1978). The conversational hand gestures of deaf and hearing adolescents. Journal of Communication, 28(3), 108–115.

Bavelas, J. B., Chovil, N., Lawrie, D. A., & Wade, A. (1992). Interactive gestures. Discourse Processes, 15(4), 469–489.

Bergmann, K., & Kopp, S. (2006). Verbal or visual? How information is distributed across speech and gesture in spatial dialogue. In Proceedings of the 10th Workshop on the Semantics and Pragmatics of Dialogue.

Butterworth, B., & Beattie, G. W. (1978). Gesture and silence as indicators of planning in speech. In R. Campbell & P. T. Smith (Eds.), Recent Advances in the Psychology of Language (pp. 347–360). Springer.

Butterworth, B., & Hadar, U. (1989). Gesture, speech, and computational stages: A reply to McNeill. Psychological Review, 96(1), 168–174.

Cassell, J., McNeill, D., & McCullough, K. E. (1999). Speech-gesture mismatches: Evidence for one underlying representation of linguistic and nonlinguistic information. Pragmatics and Cognition, 7(1), 1–34.

Chang, D. H. F., Troje, N. F., Ikegaya, Y., Fujita, I., & Ban, H. (2021). Spatiotemporal dynamics of responses to biological motion in the human brain. Cortex; a Journal Devoted to the Study of the Nervous System and Behavior, 136, 124–139. 10.1016/j.cortex.2020.12.015

De Ruiter, J. P. (1998). Gesture and Speech Production. Unpublished Doctoral Dissertation. University of Nijmegen.

De Ruiter, J. P. (2000). The production of gesture and speech. In D. McNeill (Ed.), Language and Gesture (pp. 284–311). Cambridge University Press.

Dick, A. S., Goldin-Meadow, S., Solodkin, A., & Small, S. L. (2008). Gesture in the developing brain. Developmental Science, 11(2), 202–213.

Dick, A. S., Goldin-Meadow, S., Hasson, U., Skipper, J., & Small, S. L. (2009). Co-speech gestures influence neural responses in brain regions associated with semantic processing. Human Brain Mapping, 30(11), 3509–3526.

Dick, A. S., Goldin-Meadow, S., Solodkin, A., & Small, S. L. (2012). Gesture in the developing brain. Developmental Science, 15(2), 165–180. 10.1111/j.1467-7687.2011.01100.x

Dittmann, A. T., & Llewellyn, L. G. (1969). Relationship between vocal pauses and gestures. Journal of Personality and Social Psychology, 11(6), 498–504.

Dobrogaev, S. V. (1929). *Gestures and speech: An experimental study*. In related-potentials (pp. 263–297). Cambridge, MA: The MIT Press.

Demir-Lira, Ö. E., Asaridou, S. S., Raja Beharelle, A., Holt, A. E., Goldin-Meadow, S., & Small, S. L. (2018). Functional neuroanatomy of gesture-speech integration in children varies with individual differences in gesture processing. Developmental Science, 21(5), e12648.

Drijvers, L., & Özyürek, A. (2018). Native language status of the listener modulates the neural integration of speech and iconic gestures in clear and adverse listening conditions. Brain and Language, 177-178, 7–17.

Graham, J. A., & Heywood, S. (1975). The effects of elimination of hand gestures and of verbal codability on speech performance. European Journal of Social Psychology, 5(2), 189–195. 10.1002/ejsp.2420050204

Gramfort, A., Luessi, M., Larson, E., Engemann, D. A., Strohmeier, D., Brodbeck, C., Goj, R., Jas, M., Brooks, T., Parkkonen, L., & Hämäläinen, M. (2013). MEG and EEG data analysis with MNE-Python. Frontiers in Neuroscience, 7, 267. 10.3389/fnins.2013.00267

Green, A., Straube, B., Weis, S., Jansen, A., Willmes, K., Konrad, K., & Kircher, T. (2009). Neural integration of iconic and unrelated coverbal gestures: A functional MRI study. Human Brain Mapping, 30, 3309–3324.

Habets, B., Kita, S., Shao, Z., Özyürek, A., & Hagoort, P. (2011). The role of synchrony and ambiguity in speech–gesture integration during comprehension. Journal of Cognitive Neuroscience, 23(8), 1845–1854.

Hadar, U., & Butterworth, B. (1997). Iconic gestures, imagery, and word retrieval in speech. Semiotica, 115(1–2), 147–172.

Hoetjes, M., Krahmer, E., & Swerts, M. (2014). Does our speech change when we cannot gesture? Speech Communication, 57, 257–267. 10.1016/j.specom.2013.06.007

Holle, H., & Gunter, T. C. (2007). The role of iconic gestures in speech disambiguation: ERP evidence. Journal of Cognitive Neuroscience, 19(7), 1175–1192.

Holle, H., Gunter, T. C., Rüschemeyer, S. A., Hennenlotter, A., & Iacoboni, M. (2008). Neural correlates of the processing of co-speech gestures. NeuroImage, 39, 2010–2024.

Holle, H., Obleser, J., Rueschemeyer, S.-A., & Gunter, T. C. (2010). Integration of iconic gestures and speech in left superior temporal areas boosts speech comprehension under adverse listening conditions. NeuroImage, 49, 875–884.

Hyvärinen, A. (1999). Fast and robust fixed-point algorithms for independent component analysis. IEEE Transactions on Neural Networks, 10(3), 626–634. 10.1109/72.761722

Iverson, J. M., & Goldin-Meadow, S. (1997). What’s communication got to do with it? Gesture in blind children. Developmental Psychology, 33(3), 453–467.

Kelly, S. D., Kravitz, C., & Hopkins, M. (2004). Neural correlates of bimodal speech and gesture comprehension. Brain and Language, 89, 253–260.

Kısa, Y. D., Goldin-Meadow, S., & Casasanto, D. (2021). Do Gestures Really Facilitate Speech Production?. Journal of Experimental Psychology: General, 151(6), 1252–1271. 10.1037/xge0001135

Kita, S. (2000). How representational gestures help speaking. In D. McNeill (Ed.), Language and Gesture (pp. 162–185). Cambridge University Press.

Kita, S., & Davies, T. S. (2009). Competing conceptual representations trigger co-speech representational gestures. Language and Cognitive Processes, 24(5), 761–775. 10.1080/01690960802327971

Kita, S., & Özyürek, A. (2003). What does cross-linguistic variation in semantic coordination of speech and gesture reveal?: Evidence for an interface representation of spatial thinking and speaking. Journal of Memory and Language, 48, 16–32.

Krauss, R. M., Morrel-Samuels, P., & Colasante, C. (1991). Do conversational hand gestures communicate? Journal of Personality and Social Psychology, 61, 743–759.

Krauss, R. M., Chen, Y., & Gottesman, R. F. (2000). Lexical gestures and lexical access: A process model. In D. McNeill (Ed.), Language and Gesture (pp. 261–283). Cambridge University Press.

Larson, E., & Taulu, S. (2018). Reducing sensor noise in MEG and EEG recordings using oversampled temporal projection. IEEE Transactions on Bio-Medical Engineering, 65(5), 1002–1013. 10.1109/TBME.2017.2734641

Lausberg, H., & Sloetjes, H. (2009). Coding gestural behavior with the NEUROGES-ELAN system. Behavior Research Methods, 41(3), 841–849. Retrieved from http://tla.mpi.nl/tools/tla-tools/elan/

Lambon Ralph, M. A., Jefferies, E., Patterson, K., & Rogers, T. T. (2017). The neural and computational bases of semantic cognition. Nature Reviews Neuroscience, 18(1), 42–55.

Levelt, W. J. M. (1989). Speaking: From intention to articulation. MIT Press.

Mamus, E., Speed, L. J., Rissman, L., Majid, A., & Özyürek, A. (2023). Lack of visual experience affects multimodal language production: Evidence from congenitally blind and sighted people. Cognitive Science, 47(1), e13228. 10.1111/cogs.13228

McNeill, D. (1992). Hand and mind: What gestures reveal about thought. University of Chicago Press.

McNeill, D. (2005). Gesture and thought. University of Chicago Press.

Nobe, S. (2000). Where do most spontaneous representational gestures actually occur with respect to speech? In D. McNeill (Ed.), Language and gesture (pp. 186–198). Cambridge University Press.

Oi, M., Saito, H., Li, Z., & Zhao, W. (2013). Co-speech gesture production in an animation–narration task by bilinguals: A near-infrared spectroscopy study. Brain and Language, 125(1), 77–81. 10.1016/j.bandl.2013.01.004

Özyürek, A. (2014). Hearing and seeing meaning in speech and gesture: Insights from brain and behaviour. Philosophical Transactions of the Royal Society of London, 369. 10.1098/rstb.2013.0296

Özyürek, A., Willems, R. M., Kita, S., & Hagoort, P. (2007). On-line integration of semantic information from speech and gesture: Insights from event-related brain potentials. Journal of Cognitive Neuroscience, 19, 605–616.

Patterson, K., Nestor, P. J., & Rogers, T. T. (2007). Where do you know what you know? The representation of semantic knowledge in the human brain. Nature Reviews Neuroscience, 8(12), 976–987.

Rauscher, F. H., Krauss, R. M., & Chen, Y. (1996). Gesture, speech, and lexical access: The role of lexical movements in speech production. Psychological Science, 7(4), 226–231.

Rimé, B., Schiaratura, L., Hupet, M., & Ghysselinckx, A. (1984). Effects of relative immobilization on the speaker’s nonverbal behavior and on the dialogue imagery level. Motivation and Emotion, 8, 311–325.

Sato, Y., Nishimaru, H., Matsumoto, J., Setogawa, T., & Nishijo, H. (2023). Electroencephalographic effective connectivity analysis of the neural networks during gesture and speech production planning in young adults. Brain Sciences, 13(1), 100. 10.3390/brainsci13010100

Sekine, K., Sowden, H., & Kita, S. (2015). The development of the ability to semantically integrate information in speech and iconic gesture in comprehension. Cognitive Science, 39, 1855–1888.

Sekine, K., Schoechl, C., Mulder, K., Holler, J., Kelly, S., Furman, R., & Özyürek, A. (2020). Evidence for children’s online integration of simultaneous information from speech and iconic gestures: An ERP study. *Language*, Cognition and Neuroscience, 35(10), 1283–1291. 10.1080/23273798.2020.1737719

Straube, B., Green, A., Weis, S., & Chatterjee, A. (2009). Memory effects of speech and gesture binding: Cortical and hippocampal activation in relation to subsequent memory performance. Journal of Cognitive Neuroscience, 21, 821–836.

Taulu, S., Simola, J., & Kajola, M. (2005). Applications of the signal space separation method. IEEE Transactions on Signal Processing: A Publication of the IEEE Signal Processing Society, 53(9), 3359–3372. 10.1109/TSP.2005.853302

Theochaaropoulou, F., Cocks, N., Pring, T. and Dipper, L. (2015). “TOT” phenomena: Gesture production in younger and older adults. Psychology and Aging, 30(2), 245–252. doi: 10.1037/a0038913

Wang, X., Lu, K., He, Y. et al. Dynamic brain networks in spontaneous gestural communication. npj Science of Learning, 9, 59 (2024). 10.1038/s41539-024-00274-2

Willems, R. M., Özyürek, A., & Hagoort, P. (2008). Seeing and hearing meaning: ERP and fMRI evidence of word versus picture integration into a sentence context. Journal of Cognitive Neuroscience, 20, 1235–1249.

