## Supplemental_figures for "The time course of co-speech gesture production: An MEG study"

**Supplementary Materials**

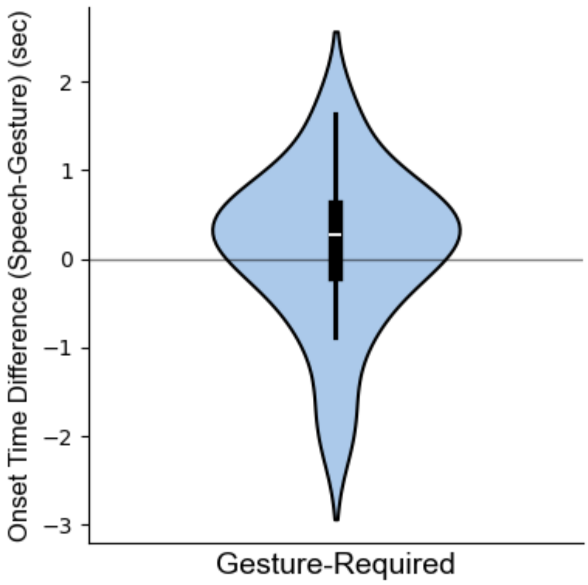


**Figure S1. Difference in onset time between speech and gesture**

Only the Gesture-Required condition was analyzed. Gestures did not consistently occur either before or after speech onset, M ± SD = 0.114 ± 0.849 sec. A one-sample t-test against zero showed no significant difference, $t(22) = 0.632$, $p = 0.534$.

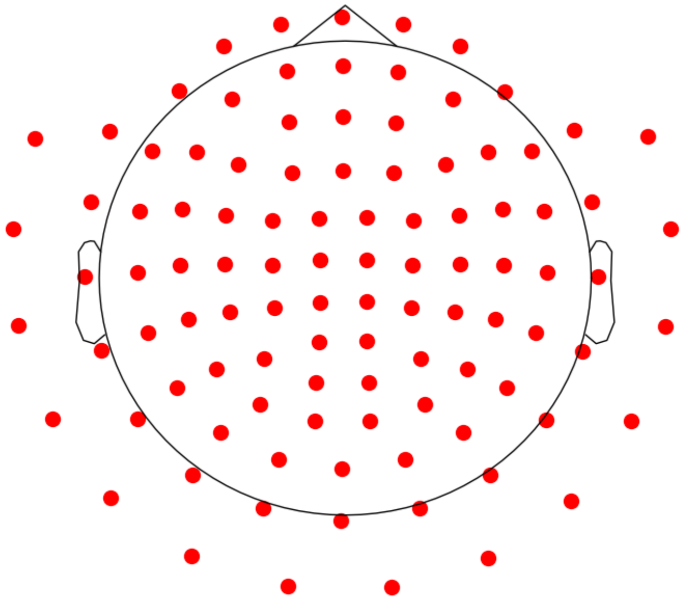

**Figure S2. Layout of MEG channels**

Of the 360 measured channels, 102 magnetometer channels were used for analysis.

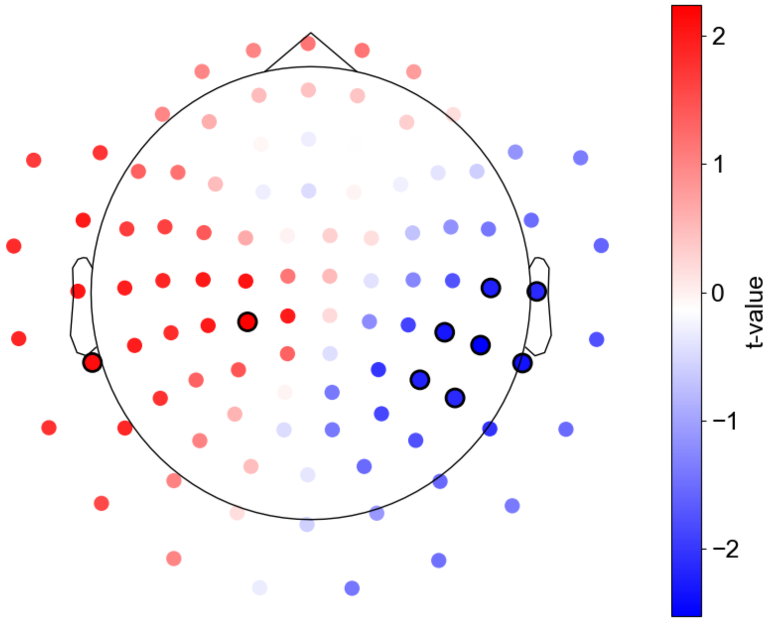


**Figure S3. Channels showing differences in mean amplitude (–0.25 to 0 sec relative to speech onset)**

To identify any differences outside the bilateral ATL, we conducted paired t-tests on the mean amplitude of each channel. Channels with uncorrected p-values < 0.05 are marked with black circles.

No significant differences were found beyond the ATL.

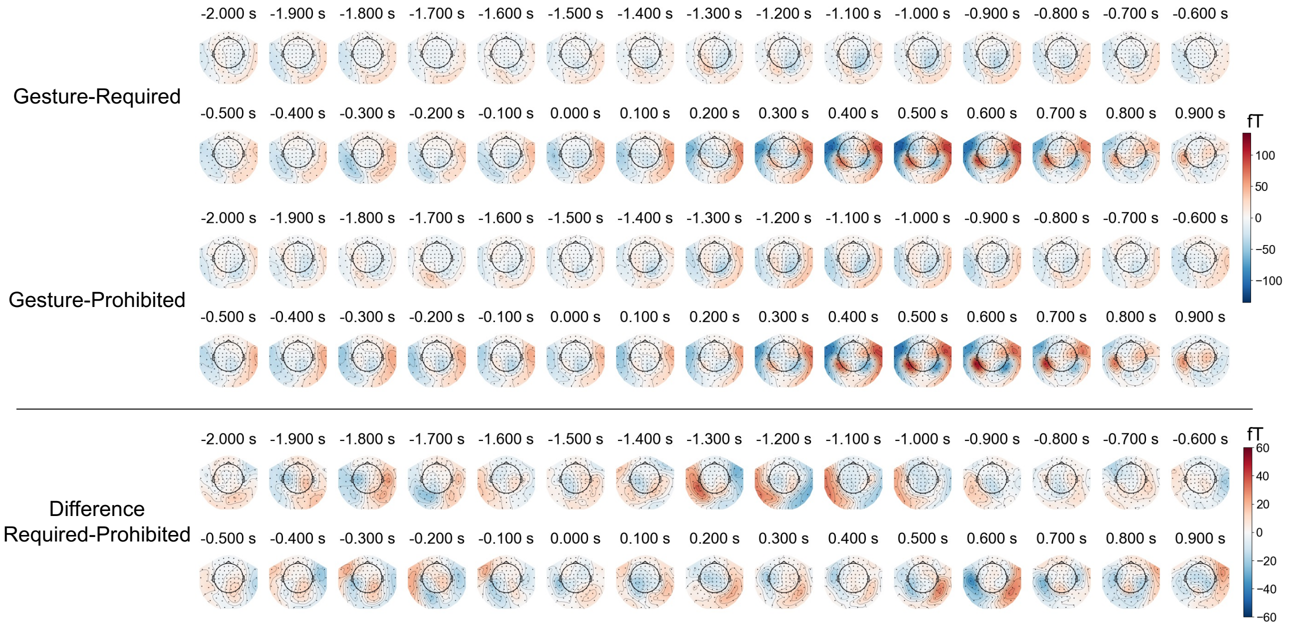


**Figure S4. Time series of topomaps around trial onset (-2 to 1 sec)**

Time 0 indicates the trial onset. No differences were observed between the conditions. Top: Gesture-Required condition; Middle: Gesture-Prohibited condition; Bottom: Difference between the two conditions.

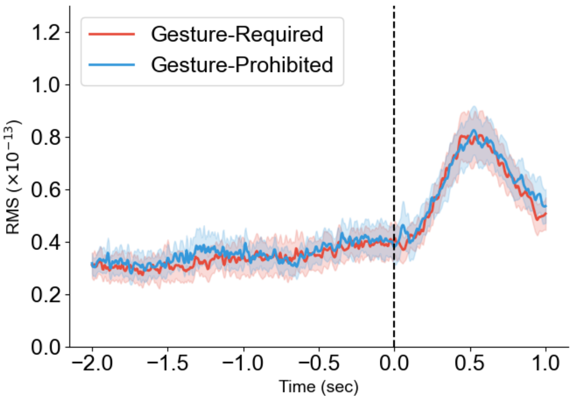


**Figure S5. Root mean square (RMS) aligned to trial onset**

No difference was observed between the Gesture-Required and Gesture-Prohibited conditions. No statistical tests were conducted.

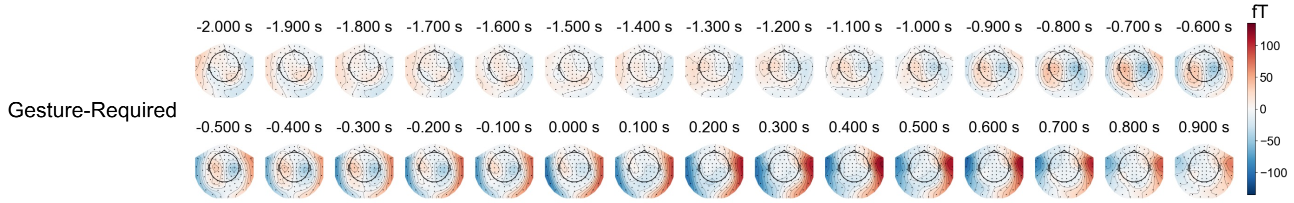


**Figure S6. Time series of topomaps around gesture onset (-2 to 1 sec)**

Time 0 indicates the gesture onset. Activity in the motor area was observed starting approximately 0.3 s before gesture onset.
